# Multi-timescale hybrid components of the functional brain connectome: A bimodal EEG-fMRI decomposition

**DOI:** 10.1101/800797

**Authors:** Jonathan Wirsich, Enrico Amico, Anne-Lise Giraud, Joaquín Goñi, Sepideh Sadaghiani

## Abstract

Concurrent electroencephalography (EEG) and functional magnetic resonance imaging (fMRI) bridge brain connectivity across timescales. During concurrent EEG-fMRI resting-state recordings, whole-brain functional connectivity (FC) strength is spatially correlated across modalities. However, cross-modal investigations have commonly remained correlational, and joint analysis of EEG-fMRI connectivity is largely unexplored.

Here we investigated if there exist (spatially) independent FC networks linked between modalities. We applied the recently proposed hybrid connectivity independent component analysis (connICA) framework to two concurrent EEG-fMRI resting state datasets (total 40 subjects). Two robust components were found across both datasets. The first component has a uniformly distributed EEG-frequency fingerprint linked mainly to intrinsic connectivity networks (ICNs) in both modalities. Conversely, the second component is sensitive to different EEG-frequencies and is primarily linked to intra-ICN connectivity in fMRI but to inter-ICN connectivity in EEG.

The first hybrid component suggests that connectivity dynamics within well-known ICNs span timescales, from millisecond-range in all canonical frequencies of FC_EEG_ to second-range of FC_fMRI_. Conversely, the second component additionally exposes linked but spatially divergent neuronal processing at the two timescales. This work reveals the existence of joint spatially independent components, suggesting that parts of resting-state connectivity are co-expressed in a linked manner across EEG and fMRI over individuals.

## Introduction

The advances in neuroimaging in the last decades have brought new important insights on brain functioning using both electroencephalography (EEG) data and functional magnetic resonance imaging (fMRI) data extracted from human brain. EEG provides a direct measure of neuronal activity with high temporal resolution but is limited by the spatial coverage of electrodes on the scalp and by volume conductance. Conversely, fMRI provides a high-resolution estimate of brain function with whole-brain coverage, however with limited temporal reliability restricted by the slow filter of the hemodynamic response (Logothetis et al., 2001) and the time needed to acquire a whole brain image (typically in the range of 1-3 seconds).

Functional connectivity, measured as statistical dependence of neural activity across distant brain areas, is considered crucial for brain function and cognition. In large-scale brain network models of functional connectivity (FC), nodes correspond to gray-matter regions (based on brain atlases or parcellations) while links or edges correspond to connections between the nodes. The connectivity between brain regions in fMRI is usually computed as the pairwise Pearson’s correlation between brain regions’ infraslow (<0.1 Hz) time series (Biswal et al., 1995). On a whole-brain level this graph is either called a functional connectivity matrix (Achard et al., 2006) or functional connectome (FC_fMRI_) (Sporns, 2011). When looking at the whole brain functional connectivity of the human brain a stable pattern of inter-connected intrinsic connectivity network (ICNs) arise that closely resemble co-activation patterns observed during cognitive tasks (Damoiseaux et al., 2006; Fox et al., 2005; Yeo et al., 2011).

In electrophysiological recordings, stable activation patterns arise forming so-called microstates (Koukkou-Lehmann et al., 1980; Lehmann et al., 1987) reorganizing on a fast timescale (around 100ms). It has been shown that these fast electrophysiological patterns are linked to fMRI ICNs (Britz et al., 2010; Musso et al., 2010). Beyond such activation patterns, to derive electrophysiological *connectivity* in MEG or EEG (FC_M/EEG_), the statistical dependencies between brain regions’ M/EEG activity can be measured by phase coupling e.g. using the pairwise imaginary part of the coherency (Wirsich et al., 2017) or by amplitude coupling using Pearson’s correlation of the Hilbert envelope (Brookes et al., 2011; Deligianni et al., 2014). These pairwise FC_M/EEG_ measures are estimated for source-reconstructed M/EEG time series at a specific frequency band. In other words, the rich oscillatory repertoire of electrophysiological activity gives rise to multiple frequency-specific connectomes in various canonical frequency bands. Phase- and amplitude-based connectivity measures are reliably related to each other (Colclough et al., 2016), and it has been shown that both organize into ICNs (de Pasquale et al., 2010; de Pasquale et al., 2012) spatially resembling fMRI ICNs (Brookes et al., 2011). Recent concurrent EEG-fMRI studies have directly shown that whole-brain EEG connectivity in all oscillatory frequency bands is spatially related to fMRI connectivity (Deligianni et al. 2014, Wirsich et al. 2017).

Besides this partial connectivity overlap across modalities, brain responses measured from fMRI and electrophysiology show discrepancies not only arising from different signal-to-noise levels, but also because they capture different aspects of brain activity at different timescales (Furey et al., 2006). Those discrepancies can also be observed when comparing multimodal connectivity. Indeed, when compared directly, it was recently shown that modality-specific differences in localized pairwise connections between FC_fMRI_ and FC_EEG_ when relating each of the functional modalities to diffusion MRI (dMRI) connectivity (Wirsich et al., 2017). Specifically, when trying to explain the underlying structural connectivity using the combination of fMRI- and EEG-derived functional connectivity, we observed that EEG-delta connectivity provides additional information to fMRI connectivity at the global level. Conversely, gamma contributes information beyond FC_fMRI_ locally in areas of the visual cortex. To date, the few existing investigations of concurrently recorded fMRI and source-space EEG connectomes (Deligianni et al., 2016, 2014; Wirsich et al., 2017) remain correlational, and analysis of joint EEG-fMRI connectivity is largely unexplored. In consequence, the relationship between EEG and fMRI connectivity organization is still incompletely characterized.

Recently Amico et al. (2017) have proposed applying independent component analysis (ICA) to whole brain functional connectivity to extract independent components, also termed as independent connectivity traits. This connectivity ICA (connICA) framework has been shown to be useful in extracting the hybrid independent components jointly expressed across fMRI- and diffusion MRI (dMRI)-derived connectivity (Amico and Goñi, 2018a). An interesting feature of this approach is that it is able to identify hybrid joint connectivity components that are linked in terms of explaining subject-specific variance of spatially independent non-linear relationships between both modalities. Note that in contrast to other bimodal ICA approaches (e.g. (Eichele et al., 2009)), hybrid connICA operates on connectivity values rather than on signal magnitudes or their timeseries. As such, the connICA approach provides the optimal framework to access multi-timescale connectome components. This is the case not only if the modalities contribute in a spatially consistent manner but also in a spatially divergent manner to the variance observed in different subjects.

Firstly, we asked if there exist spatially independent brain network patterns co-occurring over modalities in simultaneous EEG-fMRI data. Secondly, we were interested in determining how many independent connectivity patterns are collapsed into the global connectivity organization provided by FCs_fMRI_ and FCs_EEG_. And thirdly, we sought to determine the contribution of different oscillation frequencies to the EEG connectivity organization, and how this frequency distribution relates to fMRI connectivity.

In summary, in this study we sought to answer if there exist spatially independent frequency-specific FC patterns co-expressed between modalities across subjects by extending the hybrid connICA approach proposed in (Amico and Goñi, 2018a) to concurrent EEG-fMRI resting-state data.

## Methods

We use two independent datasets acquired from two different sites. The main dataset consisted of 26 healthy subjects with 3 runs of 10 minutes of concurrent EEG-fMRI during task-free resting-state (Sadaghiani et al., 2010). The generalization dataset consisted of 14 subjects, 20 minutes resting-state runs of concurrent EEG-fMRI (Wirsich et al., 2017). The data was used to extract the main joint independent components co-occurring during EEG-fMRI resting-state.

### Main Dataset

MR was acquired in 26 healthy subjects (8 females, mean age 24.39, age range 18-31) with no history of neurological or psychiatric illness. Ethical approval has been obtained from the local Research Ethics Committee (Comité de Protection des Personnes (CPP) Ile de France III) and informed consent has been obtained from all subjects.

Three runs of 10 minutes eyes-closed resting-state were acquired in one concurrent EEG-fMRI session (Tim-Trio 3T, Siemens, 40 slices, TR=2.0s, 3.0×3.0×3.0mm, TE = 50ms, field of view 192, FA=78°). EEG was simultaneously recorded using an MR-compatible amplifier (BrainAmp MR, sampling rate 5kHz), 62 electrodes (Easycap), referenced to FCz, 1 ECG electrode, and 1 EOG electrode, while the scanner clock was time-locked with the amplifier clock (Mandelkow et al., 2006). An anatomical T1-weighted MPRAGE (176 slices, 1.0×1.0×1.0 mm, field of view 256, TR=7min) was equally acquired. The acquisition was part of a study with two naturalistic film stimuli of 10 minutes not analyzed in this study (acquired after runs 1 and 2 of the resting state as described in Morillon et al. (2010)). Subjects wore earplugs to attenuate scanner noise and were asked to stay awake, avoid movement and close their eyes during resting-state recordings. Due to insufficient EEG quality in three subjects, one of three rest sessions were excluded.

#### Brain parcellation

T1-weighted images were processed with the Freesurfer suite (recon-all, v6.0, http://surfer.nmr.mgh.harvard.edu/) performing non-uniformity and intensity correction, skull stripping and grey/white matter segmentation. The cortex was parcellated into 148 regions according to the Destrieux atlas (Destrieux et al., 2010; Fischl et al., 2004).

#### fMRI processing

The fMRI timeseries were subjected to timeslicing followed by spatial realignment using the SPM12 toolbox (revision 6906, http://www.fil.ion.ucl.ac.uk/spm/software/spm12). The subjects’ T1 image and Destrieux atlas were coregistered to the fMRI images. Average CSF and white matter signal from manually defined ROIs (5mm sphere, Marsbar Toolbox 0.44, http://marsbar.sourceforge.net) were extracted and were regressed out of the BOLD timeseries along with 6 rotation and translation motion parameters and global gray matter signal. Timeseries were bandpass-filtered at 0.009-0.08 Hz (Power et al., 2014) and scrubbed using frame wise displacement (threshold 0.5) defined by Power et al. (Power et al., 2012).

#### EEG processing

The gradient artefact induced by the scanner on the EEG signal was removed using the template subtraction and adaptive noise cancelation followed by lowpass filtering at 75Hz, downsampling to 250Hz (Allen et al., 1998). Then a cardiobalistic artefact template subtraction (Allen et al., 2000) was carried out, using EEGlab v.7 (http://sccn.ucsd.edu/eeglab) and the FMRIB plug-in (https://fsl.fmrib.ox.ac.uk/eeglab/fmribplugin/). Data was then analyzed with Brainstorm software (Tadel et al., 2011), which is documented and freely available under the GNU general public license (http://neuroimage.usc.edu/brainstorm, version August 2017). Data was bandpass-filtered at 0.3-70 Hz and segmented according to the TR of the fMRI acquisition (2s epochs). Epochs that contained head motion artifacts in EEG were visually identified after semi-automatically preselecting epochs where signal in any channel exceeded the mean channel timecourse by 4 std.

Electrode positions were manually co-registered to the T1 image and a forward model of the skull was calculated using the T1 image of each subject using the OpenMEEG BEM model (Gramfort et al., 2010; Kybic et al., 2005).

EEG was re-referenced to the global average, and data were reconstructed into source space using the Tikhonov-regularized minimum-norm with the Tikhonov parameter set to λ = 10% of maximum singular value of the lead field (Baillet et al., 2001). Source timeseries were averaged to the regions of the Destrieux atlas and connectivity matrices were calculated for each segment by taking the imaginary coherency of delta, theta, alpha, beta and gamma frequency bands between each region. Excluding the real part of coherency is a common method to avoid spurious connectivity stemming from volume conductance (Nolte et al., 2004). The final EEG connectivity matrices were obtained by averaging the band specific connectivity of all segments (FC_δ_, FC_θ_, FC_α_, FC_β_ and FC_γ_). All steps of EEG and fMRI connectome construction are summarized in Figure 1.

**Figure 1:**
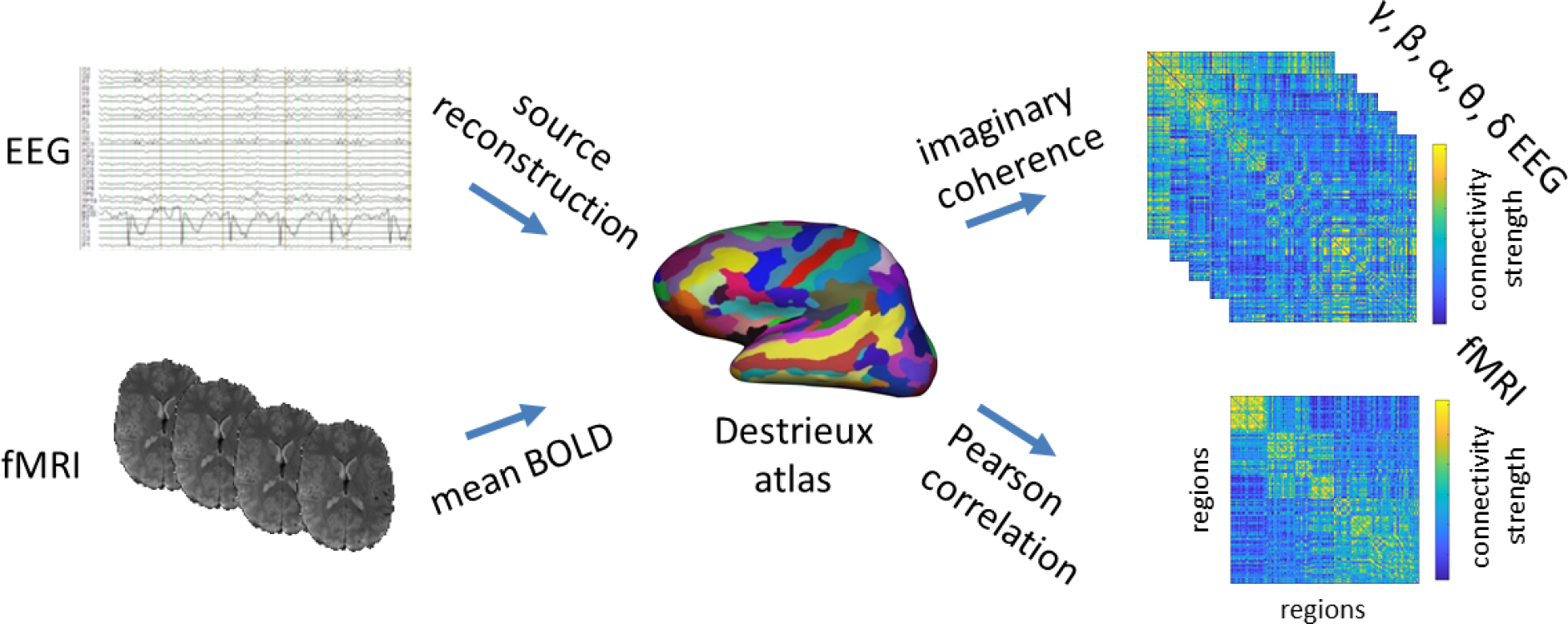
Construction of EEG and fMRI connectomes. EEG and fMRI data were parcellated into the 148 cortical regions of the Destrieux atlas, Pearsons correlation of fMRI timecourses and imaginary part of the coherency of EEG source signal were use to build connectomes.

#### Independent component analysis

We used the hybrid connectivity independent component analysis (connICA, (Amico et al., 2017; Amico and Goñi, 2018a), code available here: https://engineering.purdue.edu/ConnplexityLab/publications/connICA_hybrid_toolbox_v1.0.tar.gz) method as a data-driven way to disentangle the main brain network patterns underlying the FCs_fMRI_ and FCs_EEG_ (Figure 2).

**Figure 2:**
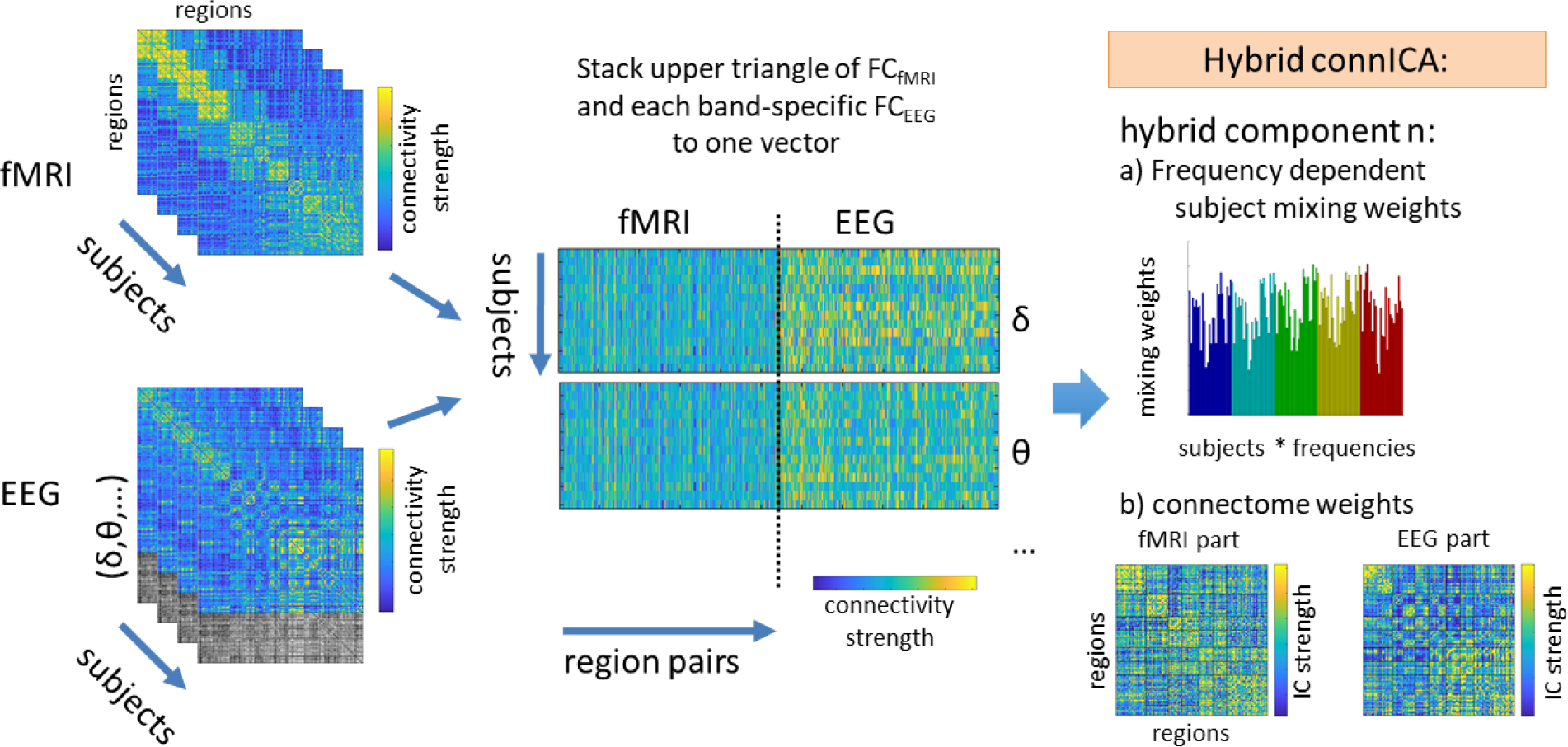
The upper triangular of each individual fMRI functional connectivity (FC_fMRI_) matrix (left, top) and lower triangular of each correspondent EEG functional connectivity (FC_EEG_) matrix (left, bottom) are added to a matrix where rows are the subjects*EEG frequency bands and columns are their vectorized hybrid (fMRI-EEG) connectivity patterns, the fMRI vector is repeated and concatenated with each of the five canonical frequency bands (middle). The ICA algorithm extracts the n independent components (i.e. ICs) associated to the whole population and their relative ICA mixing weights across subjects*EEG frequency bands (right).

To analyze the data in one hybrid vector, we uniformed the value distribution range of EEG (imaginary coherency original range 0 to 1) and fMRI (Pearson correlation, range −1 to 1) by transforming the EEG imaginary coherency according to the procedure in (Amico and Goñi, 2018a). In brief, we derived a correlation value from EEG by taking the Pearson correlation between the i_th_ and j_th_ row of the imaginary coherency-based connectome. This procedure results in a new correlation matrix where the values −1 and 1 indicate if the EEG nodes are connected antagonistically to the network or have high similarity (see also matching index in (Rubinov and Sporns, 2010)).

Then, for each subject, we first computed the concurrent FCs_fMRI_ and FCs_EEG_ at different frequency bands (δ,ϑ,α,β,γ). We then stacked the upper and lower triangular parts of each individual FC_fMRI_ and FC_EEG_ and added it to a matrix where rows are the subjects and columns are their full connectivity pattern (see scheme in Fig.1). Second, we applied a PCA dimensionality reduction followed by an ICA run. As both PCA reduction and number of selected ICs are an arbitrary choice, parameters were explored for keeping the PCs that explain 75% to 90% of the data (in 5% steps) and reducing the data from 5 to 20 ICs (in steps of 5). The ICA algorithm was run (500 times, similarly to (Amico and Goñi, 2018a)) to extract the main hybrid (joint FC_EEG_-FC_fMRI_) components associated with the whole population. Third, the most robust components (appearing at least 75% of the times with correlation higher than 0.75 of both connectivity IC strength and subject mixing weights, see (Amico et al., 2017)) were selected.

Two consistent hybrid independent components were observed across the majority of parameter pairs (the above described PCA reduction followed by limiting the number of ICs) and across both datasets. We chose the parameters by maximizing the correlation between both datasets of those two ICs (SI Figure 1,2,3). Choosing different parameters for each dataset can result in slightly higher correlation of ICs across datasets (i.e. keeping PCs that explain 75% of the variance in one dataset and 80% of the variance in the other, SI Figure 3). In order to avoid overfitting, we chose the same configuration for both main and generalization datasets. For some parameter pairs we also observed that in the main dataset stable components sometimes split up into two components. Both components are correlated to one component in the generalization dataset and can optionally be merged into one component that correlates even better with the component found in the generalization data. In case we found two components in the main dataset we kept only the component that correlated more with the generalization dataset (to conservatively avoid adding an additional merging step).

Finally we kept the IC configuration most consistent between both data sets: PCA was applied and the first components explaining 75% of the variance (12 first PCs in case of the main dataset and 14 first PCs for the generalization dataset) were kept for further analysis.

#### ICC of mixing weights

In order to characterize the properties of the resulting hybrid ICs we assessed Graph modularity and Intra-class correlation (ICC (Bartko, 1966)) of both EEG frequency-specific and subject-specific mixing weights.

#### Modularity of ICs

Modularity of EEG and fMRI IC matrix was calculated based on the Brain Connectivity Toolbox (version 2019_03_03, https://sites.google.com/site/bctnet/, (Rubinov and Sporns, 2010)). To test in how far the networks overlap with canonical ICNs (as proposed by (Amico et al., 2017)) we calculated modularity based on a predefined community label for each region derived from the Yeo7 ICN networks (Yeo et al., 2011). For the calculation of modularity negative and positive connectivity values were treated separately. Connectivity values were treated according to the negative asymmetry implementation proposed by Rubinov and Sporns (2011) weighting positive connections more than negative connections (see function community_louvain.m in the Brain Connectivity Toolbox).

### Generalization Dataset

MR was acquired in 14 healthy subjects (five females, mean age: 30.9, std: 8,6, Min/Max age: 20-55) with no history of neurological or psychiatric illness (see (Wirsich et al., 2017)). Ethical approval has been obtained from the local Research Ethics Committee (CPP Marseille 2) and informed consent has been obtained from all subjects.

A session of 21 minutes eyes-closed resting-state was acquired using concurrent EEG-fMRI (Siemens Magnetom Verio 3T MRI-Scanner, 50 slices, TR=3.6, 2.0 × 2.0 × 2.5 mm, TE = 27 ms, FA = 90°, a total of 350 volumes). EEG-fMRI were acquired during eyes-closed resting state from Inside a (Siemens, Erlangen, Germany) subjects were wearing a 64 channel EEG-cap (BrainCap-MR 3-0, Easycap, Hersching, Germany, according to the 10–20 system with one ECG channel and a reference at the mid-frontal FCz position). EEG was simultaneously recorded using an MR-compatible amplifier (BrainAmp MR, sampling rate 5kHz), 63 electrodes (Easycap), referenced to FCz and 1 ECG electrode, while the scanner clock was time-locked with the amplifier clock (Mandelkow et al., 2006). An anatomical T1-weighted MPRAGE (TR = 1900 ms, TE = 2.19 ms, 1.0 × 1.0 × 1.0 mm, 208 slices) was equally acquired. Subjects wore noise protection to attenuate scanner noise and were asked to stay awake, avoid movement and close their eyes during resting-state recordings.

#### Data processing

Data processing was carried out as described in (Wirsich et al., 2017). In brief, the data was processed as described above (using versions Freesurfer 5.3, SPM12 6685, Marsbar 0.43). After regressing out movement, CSF, whiter matter and global gray matter signal, instead of bandpass filtering as used in the main dataset, data was filtered by wavelet analysis using the Brainwaver toolbox (version 1.6, http://cran.r-project.org/web/packages/brainwaver/index.html) (Achard et al., 2006; Wirsich et al., 2017). The Pearson correlation of the remaining wavelet coefficients of scale two (equivalent to a frequency band 0.04–0.09 Hz) was used to build a functional connectivity matrix (FC_fMRI_).

#### EEG processing

EEG artefact correction was carried out using the Brain Vision Analyzer 2 software (Brain Products, Munich, Germany). To correct for the gradient artifact a gradient template was subtracted followed by adaptive noise cancellation with 70 Hz low-pass filtering and downsampling to 250 Hz (Allen et al., 2000). Peaks from the ECG signal were extracted to generate a cardiac pulse artefact template averaged over the 100 last pulses. This template was then subtracted from the EEG signal (Allen et al., 1998). Eye movement was manually rejected using ICA and high-pass filtered at 0.3 Hz. Data was segmented into 3.6s segments (according to one TR of the fMRI sequence). Segments with obvious movement artifacts were manually excluded from the analysis. As in the main data set the remaining segments then were analyzed with Brainstorm software (Tadel et al., 2011), with the difference that used an older version of Brainstorm of January 2016 according to (Wirsich et al., 2017).

#### Independent Component Analysis, ICC and Modularity

The ICA analysis and further processing was carried out the same way as in the main dataset.

## Results

To first confirm previous correlational observations between EEG and fMRI connectivity strength on a whole brain scale (Deligianni et al., 2014; Wirsich et al., 2017), we assessed spatial correlation across the fMRI connectome and the EEG connectome in each frequency band. In line with the prior reports, correlation between group-averaged EEG and fMRI connectivity matrices were small but significant. Specifically, as reported by Wirsich et al. (2018) in the main dataset we observed correlations for fMRI vs. δ/θ/α/β/γ: r=0.34/0.34/0.33/0.36/0.29 (replication dataset correlation fMRI vs. δ/θ/α/β/γ:r=0.34/0.32/0.33/0.37/0.16 as reported in (Wirsich et al., 2017)). Next, we sought to move beyond the prior correlational approaches by using joint fMRI-EEG decomposition of brain functional connectivity.

### Determination of free parameters for connICA

When applying the connICA method to the EEG-fMRI connectivity matrices, as a function of the choice of the free parameters in the framework (PCA reduction and number of selected ICs) we observed 2-4 stable reoccurring components in the main dataset, whereas 2 stable components reoccurred in the generalization dataset using PCs explaining 75% of the variance followed by and ICA with 10 components (see methods). Two stable components were appearing in both datasets (SI Figure 1,2,3):

1. an Intrinsic coupling network’-Frequency-General component (ICN-FG) that captures all the main within-connectivity in the ICNs described by Yeo et al. (2011) (Figure 3+4). The mixing weights do not differ for different frequencies (Figure 3).
2. a Visual-Frequency-Sensitive component (VIS-FS) that shows two different patterns for fMRI and EEG that jointly co-occur. The fMRI part mainly captures the connectivity within the visual network (VIS) and the connectivity between visual and somato-motor (SM) networks (Figure 3+5), whereas the EEG part captures mainly connectivity between ICNs. Mixing weights differ according to frequency band (Figure 3).

**Figure 3:**
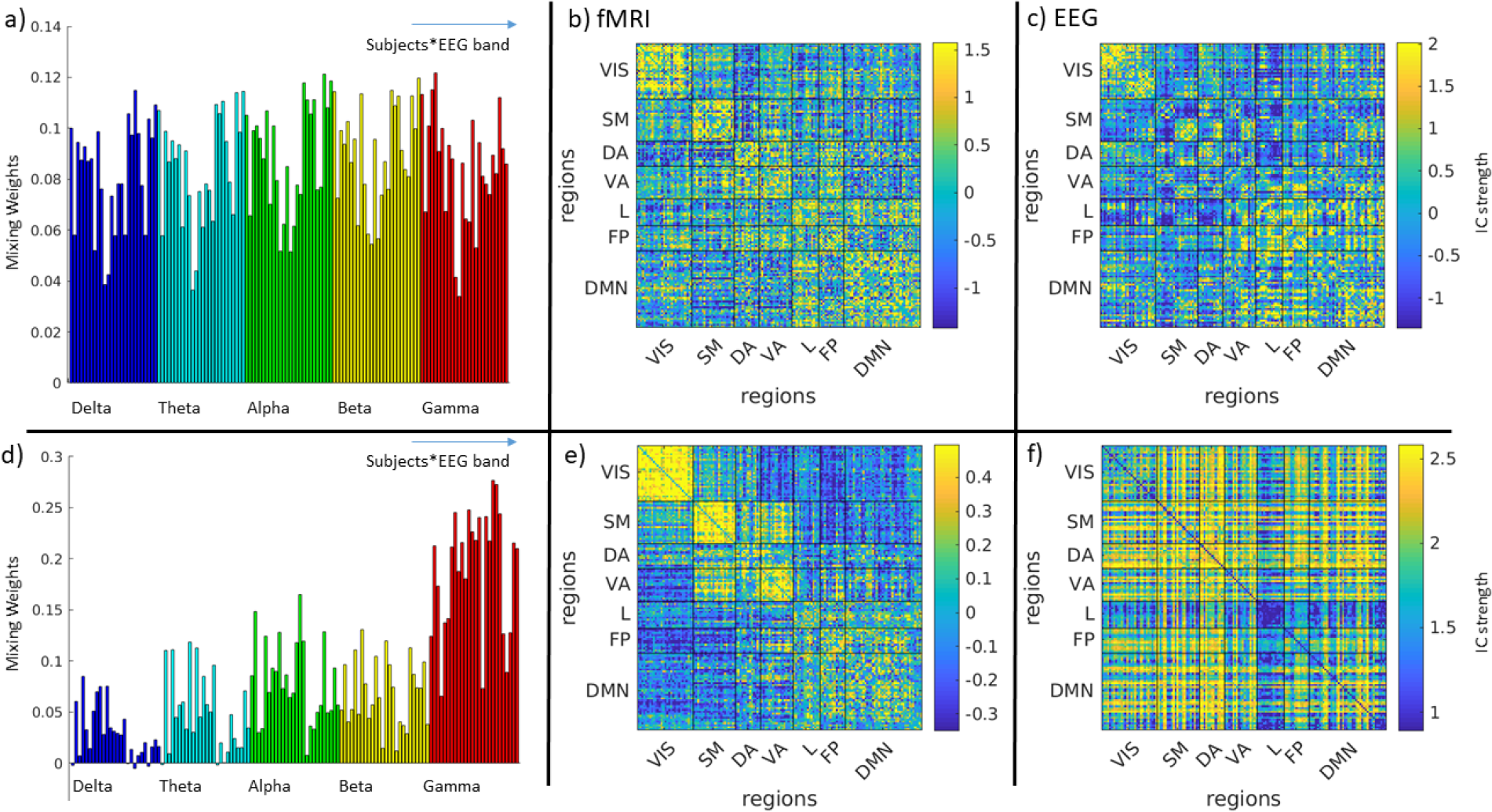
a) subject and band-specific mixing weights of the hybrid EEG-fMRI ICN-FG component b) fMRI part of the ICN-FG component c) EEG part of the ICN-FG component d) subject and band-specific IC-weights of the hybrid EEG-fMRI VIS-FS component e) fMRI part of the VIS-FS component f) EEG part of the VIS-FS component. All panels represent the main data set (for replication data see SI Figure 4, colorbars have been saturated at 95^th^ and 5^th^ percentile for better comparison with Figure SI5 and SI6). VIS: Visual, SM: Somatomotor, DA: Dorsal Attention, VA: Ventral Attention, L: Limbic, FP: Fronto Parietal, DMN: Default Mode Network

**Figure 4:**
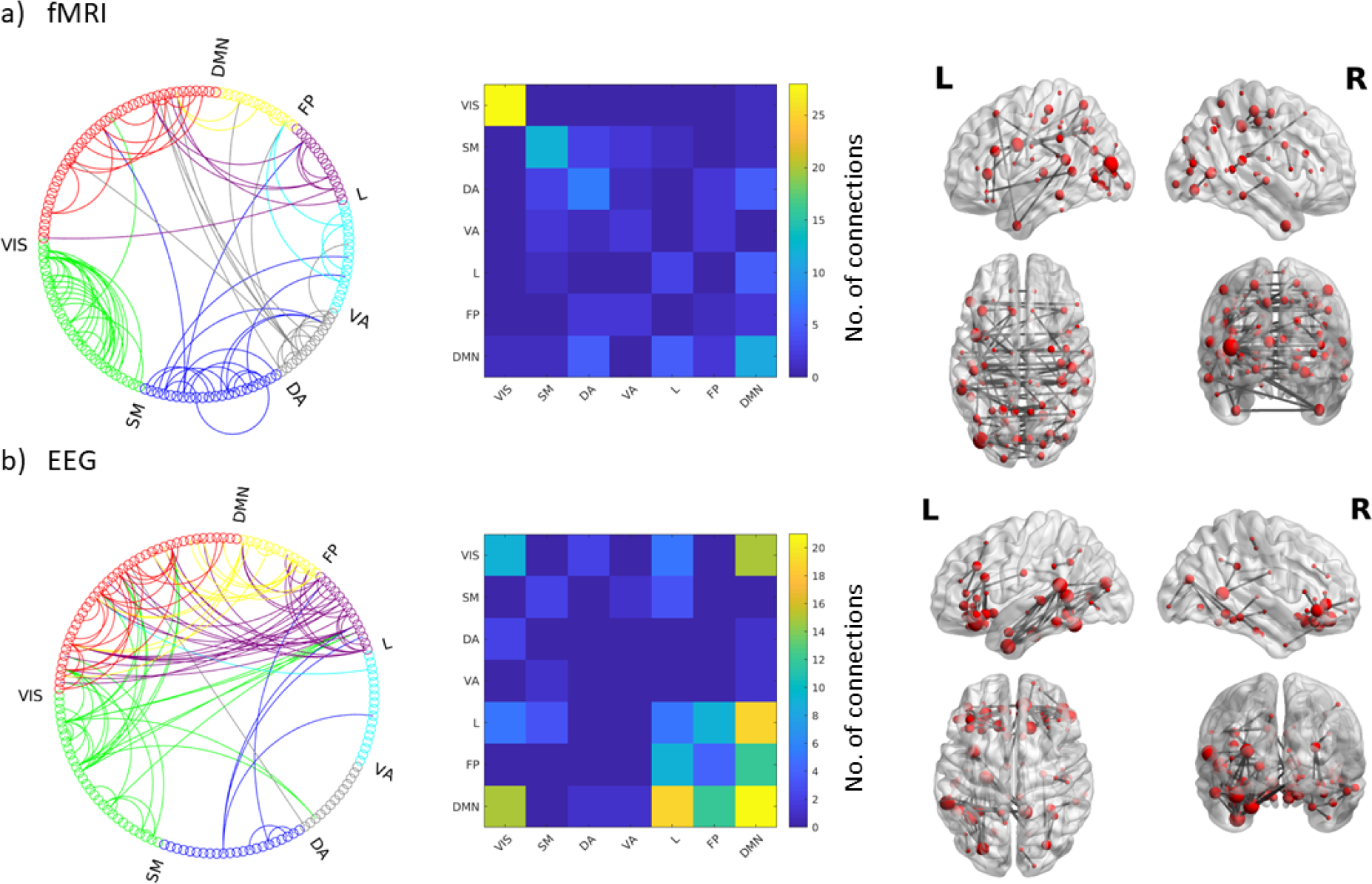
Top nodal strength connections (99^th^ percentile) for the ICN-FG component: Circle graphs show all connections between different ICN networks for fMRI (a) and EEG (d); Matrices summarize the number of connections falling into each ICN-ICN pair for fMRI (b) and EEG (e); Brain renderings show strongest connections of the components on a canonical reconstructed cortical surface for the fMRI (c) and EEG (f) part of the hyprid component. Data correspond to the main dataset; for the results if the generalization dataset see Figure SI5. VIS: Visual, SM: Somatomotor, DA: Dorsal Attention, VA: Ventral Attention, L: Limbic, FP: Fronto Parietal, DMN: Default Mode Network

**Figure 5:**
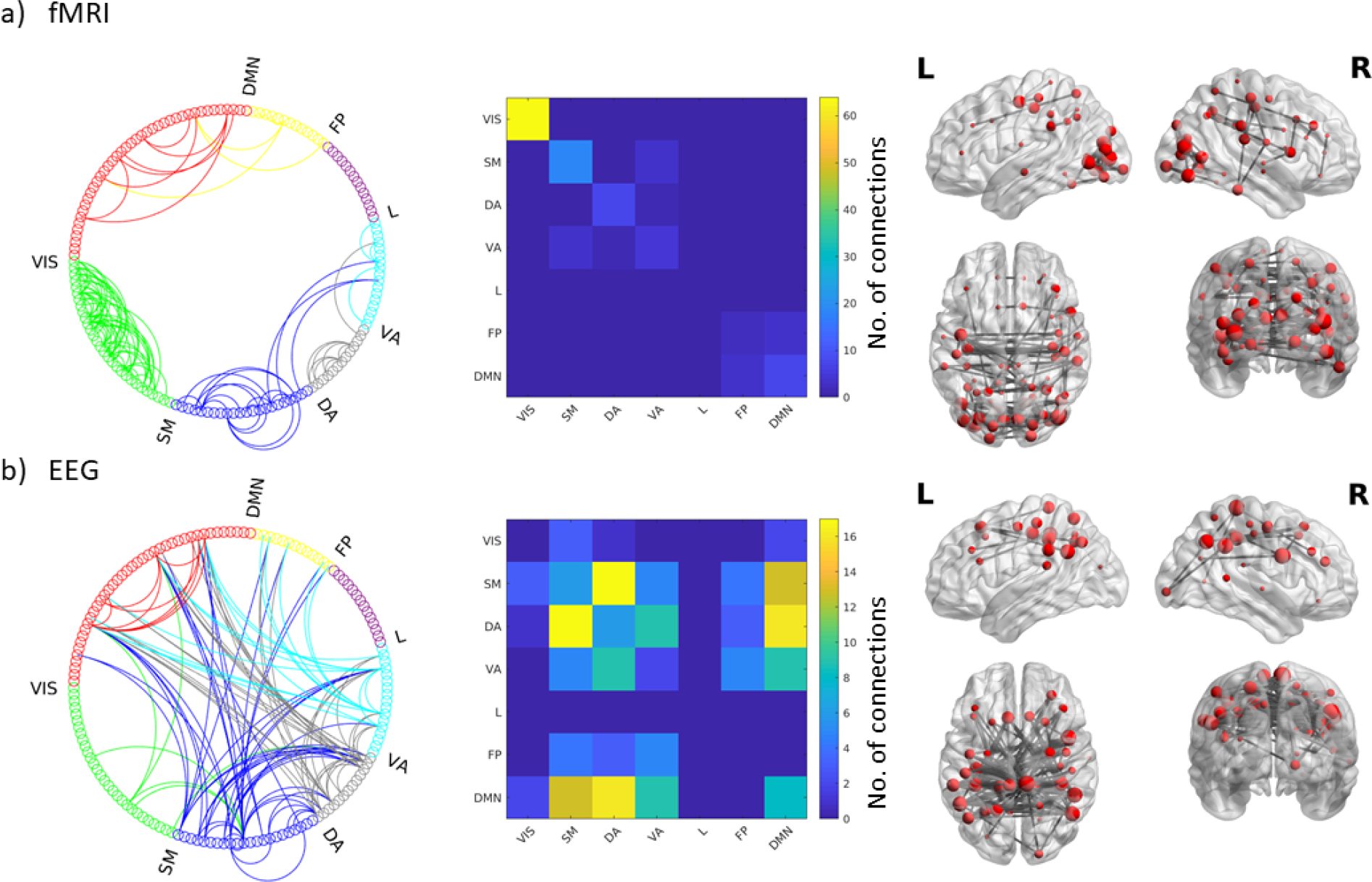
Top nodal strength connections (99^th^ percentile) for the VIS-FS component: Circle graphs show all connections between different ICN networks for fMRI (a) and EEG (d); Matrices summarize the number of connections falling into each ICN-ICN pair for fMRI (b) and EEG (e); Brain renderings show strongest connections of the components on a canonical reconstructed cortical surface for the fMRI (c) and EEG (f) part of the hyprid component. Data correspond to the main dataset; for the results if the generalization dataset see Figure SI5. VIS: Visual, SM: Somatomotor, DA: Dorsal Attention, VA: Ventral Attention, L: Limbic, FP: Fronto Parietal, DMN: Default Mode Network

### The ICN-frequency-General component

The ICN-FG component was identified in both the main and generalization datasets for both the fMRI IC weight matrix r = 0.36 (p<1.0^−300^) and the EEG IC weight matrix r=0.51 (p<1.0*10^−300^). This similarity confirms the robustness of the hybrid component across datasets.

Upon visual inspection, EEG and fMRI parts of the IC (figure 3b-c) share partial spatial similarity. This is in line with small but significant correlation between EEG and fMRI parts of the ICN-FG independent component (main data set: r=0.11, p=3.66*10^−30^; generalization data set r=0.37, p=1.42*10^−251^). Assessed visually, both EEG and fMRI parts of the IC express a strong organization into ICNs. We empirically tested this observation by calculating the modularity of the network given the predetermined labels of the 7 ICNs as defined by Yeo et al. (Yeo et al., 2011). We found a significantly high modularity as compared to and ensemble of randomized networks (5,000 iterations) with preserved weighted degree distribution and weighted degree sequence of FC_fMRI_ (Maslov and Sneppen, 2002; Rubinov and Sporns, 2011) for fMRI (q_fMRI_ = 0.22, p<.00002; generalization q_fMRI_ = 0.37, p<.00002) and moderate modularity for EEG (q_EEG_ = 0.093, p<.00002 / generalization q = 0.14, p<.00002, same null model as FC_fMRI_, 5000 iterations). The significant modularity means that connectivity is generally stronger within ICN modules than across them, confirming the relative spatial closeness of the hybrid component to the well-known ICN organization.

In addition to supporting the ICN backbone of the intrinsic connectivity architecture, the IC also showed a high subject-specific fingerprint on all its mixing weights (figure 3a, ICC_subject_=0.85, p<1.0*10^−10^; generalization ICC = 0.48, p = 2.18*10^−6^), but importantly non-significant (main data set, ICC_subject_=−0.005, p=0.42) or low (generalization data set ICC=0.27, p=0.00023673) ICC for frequency category. This result shows that the ICN-FG component has a strong contribution from all timescales and is not frequency specific.

### The Visual-Frequency-Sensitive component

The VIS-FS component was found in both datasets with r = 0.50 (p<10^−300^) for the fMRI IC matrix and r = 0.50 (p<10^−300^) for the EEG IC matrix. Again, this similarity confirms the robustness of the identified hybrid component across datasets.

Compared to the ICN-FG component, EEG and fMRI parts of the VIS-FS component diverge considerably in their spatial pattern (figure 3e-f). In line with this divergence, EEG and fMRI matrices showed a very low but significant anticorrelation (main dataset r=−0.10, p=1.09*10^−26^; generalization dataset r=−0.15, p=5.79*10^−58^). The fMRI pattern of this component is dominated by ICN organization (figure 3e) similar to the observation in the ICN-FG component. Supporting this observation, fMRI again showed significant modularity when mapped to the Yeo7 networks (compared to 5000 iterations of random networks with preserved weight and strength distribution of FC_fMRI_/FC_EEG_ (Rubinov and Sporns, 2011): q_fMRI_=0.32, p<.00002; generalization dataset q_fMRI_=0.30, p<.00002). However, the visual (VIS) ICN, and to a lesser degree the somatomotor ICN, are more strongly expressed than other ICNs. The EEG part (figure 3f) co-occurring with the aforementioned fMRI pattern depicts a more diverse connectivity profile diverging from the ICN architecture. This divergence is confirmed by lack of significant modularity when mapped to the canonical 7 ICNs (q_EEG_ −0.012, p=0.95; generalization q_EEG_=−0.018, p=0.97). The most salient characteristics of the EEG pattern are strong connectivity among default-mode network (DMN), somatomotor (SM) and dorsal attention (DA) networks (figures 3f, 4d-f).

Interestingly, these joint EEG-fMRI patterns are sensitive to different frequency bands under consideration, i.e. the mixing weights associated with the VIS-FS component change in magnitude as a function of frequency, (figure 3d, ICC = 0.69, p<1.0*10^−10^; generalization dataset ICC =0.48, p=3.51*10^−8^), and do not represent a subject-specific fingerprint (ICC = −0.11, p=0.98, generalization dataset ICC_subject_ = 0.07, p=0.19). The frequency-sensitivity was characterized by considerably stronger mixing weights for the gamma band compared to other frequencies.

### Relationship between components and head movement

To test if subject-specific mixing weights are driven by head movement, we checked for a monotonous relationship between the mixing weights and mean framewise displacement (FD), as well as between mixing weights and the number of scrubbed volumes using Spearman ranked sum correlation. For the ICN-FG component we found an negative correlation between subject-specific mixing weights and motion across all bands in the main dataset but not in the generalization dataset (main dataset: FD vs. mixing weights (figure 3a): rho = −0.44 p=1.9*10^−7^; No. of scrubbed volumes vs. ICA: rho=−0.37, p=1.4*10^−5^ / generalization data set: FD vs. mixing weights: rho=0.17, p=0.14; No. of Scrubbed Volumes vs. mixing weights: rho=0.042, p=0.73). For the VIS-FS component, we did not find a relationship between subject specific mixing weights and motion across any band in either dataset (main data set: FD vs. mixing weight (figure 3b): rho = 0.07 p=0.40; No. of scrubbed volumes vs. ICA: rho=0.12, p=0.14 / generalization data set: FD vs. mixing weights: rho=0.11, p=0.36; No. of scrubbed volumes vs. mixing weights: rho=0.04, p=0.74).

To summarize, we found a negative relationship between movement and mixing weights of the ICN-FG component in the main dataset. This outcome indicates that for the main dataset, subjects with movement contribute less to the ICN-FG component (possibly due to lower SNR in those subjects). Of note, as mentioned in the methods section we observed a second component with similar characteristics to the ICN-FG component in the main dataset, which did not show any relationship to head motion (see detailed analysis in SI Results). This observation further supports that the characteristic features of the ICN-FG component are not dependent on motion.

## Discussion

Since BOLD fMRI measures neural energy consumption, FC_fMRI_ collapses over a mixture of electrophysiological FC (FC_EEG_) mechanisms in various oscillatory frequency bands. To disentangle the complex relationship between FC_fMRI_ and multi-frequency FC_EEG_, we applied a data-driven decomposition of bimodal connectomes for simultaneous EEG-fMRI recordings. Two robust spatially independent bimodal connectome components, each representing FC_fMRI_ and FC_EEG_, were found across all frequency bands and both independent datasets. The first, that we call the Intrinsic Connectivity Network-Frequency-General component (ICN-FG), has a uniformly distributed frequency fingerprint and is linked to ICNs in both modalities (Fig. 3a, Fig. 4). Conversely, the second component that we call the Visual-Frequency-Sensitive component (VIS-FS), spatially diverges between EEG and fMRI and is sensitive to different frequency bands under consideration (i.e., the mixing weights associated to the VIS-FS component change in magnitude as a function of frequency, see Fig. 3B, Fig. 5).

### ICN-FG component

The ICN-FG component captures the main within-network connectivity in well-known neurocognitive networks for EEG and fMRI. The EEG and fMRI patterns are co-expressed over subjects and quantified by subject-specific mixing weights. This result corroborates previous findings extracting the same robust ICN-FG components from different fMRI datasets (Amico et al., 2017; Contreras et al., 2017).

The fact that we found this ICN-conform stable hybrid component across modalities might suggest that the within-network connectivity dynamics is “scale-independent”, i.e. to some extent insensitive to whether one looks at the millisecond-scale of the FC_EEG_ connectome in a specific frequency, or at the infraslow-scale of the FC_fMRI_ connectome. Previous work on microstates indeed demonstrated a temporal scale-free behavior in EEG ranging from fast milliseconds to slow second time range (Van de Ville et al., 2010). Such timescale invariance is also in line with more recent findings linking fMRI to EEG (Deligianni et al., 2014; Wirsich et al., 2017) and to MEG connectivity (Brookes et al., 2011; Colclough et al., 2016).

We found strong ICC on the subject level suggesting a strong fingerprint for each subject (Amico and Goñi, 2018b). This result should be interpreted very carefully however, as we observed in the main dataset (but not in the replication dataset) that those subject-specific mixing weights are negatively correlated with head movement for the ICN-FG component. As such, the subject-specific mixing weights should be interpreted as subject-specific contributions to the hybrid component that might also be weighted by session-specific individual contributions such as vigilance state and movement of the subject. In consequence, this also means that the final connectivity component is ‘corrected’ for those subject-specific (linear) contributions representing a clean component common among all subjects. As we showed in the main data set the ICN-FG component can split up into two parts, one being anticorrelated to movement and one being not correlated to movement. This demonstrates the ability of ICA to split up the connectivity matrix into behaviorally relevant parts depending on the number of subjects used (the larger main data set of 26 subjects was able to detect a split up of the ICN-FG component that could not be observed in the smaller generalization dataset).

In conclusion, the ICN-FG component suggests that the presence of strong electrophysiological communication in all frequencies (widely distributed, with ventral dominance) contributes to strong ICN architecture in fMRI.

### VIS-FS component

The VIS-FS hybrid component shows two spatially divergent patterns for fMRI and EEG (Figure 3c+f). The fMRI part mainly captures the connectivity within visual network (VIS) and the connectivity between visual and somato-motor (SM) networks but also other ICNs. On the other hand, the EEG part co-occurring with the aforementioned fMRI pattern depicts a more diverse connectivity profile that captures between-network connectivity especially between DMN. Though not described before, the pattern reproduces across both datasets (correlation between EEG IC strength of the two datasets r=0.50, see Fig. SI3 in SI). The fMRI IC weights of the component are in line with previous findings extracting VIS-SM independent components from different fMRI datasets (Amico et al., 2017; Contreras et al., 2017).

The observed difference across EEG and fMRI parts of the VIS-FS component suggests that the hybrid connICA is able to detect the complementary information carried over the two different timescales measured by FC_fMRI_ and FC_EEG_. As such, the γ-band-driven electrophysiological FC VIS-FS component is equally in line with the results of Wirsich et al. (2017) who observed a global contribution of delta but a local visual contribution of gamma to predicting diffusion MRI connectivity. The results reported here are consistent with this pattern as the VIS-FS component is mainly driven by gamma whereas delta does not contribute to this component and only contributes to the global pattern of the ICN-FG component (see Figure 3a).

The here observed duality of mutual and discrepant contributions of hemodynamic and electrophysiological recordings has also been discussed in task-paradigm studies. Taking face recognition tasks as an example, mutual contributions of EEG and fMRI have been readily observed (Walz et al., 2013; Wirsich et al., 2014). However, face recognition tasks also engage discrepant contributions across the two modalities; It has been shown that while the face-selective N170 originating from the fusiform gyrus in MEG is not modulated by attention, fMRI show a strong modulation in the same region (Furey et al., 2006). The divergence across modalities speaks to the hypothesis that while M/EEG weight fast conducting pathways, main contribution to the BOLD signal comes from neuronal ensembles connected via dense and slow fiber pathways (Hari and Parkkonen, 2015).

From a systems neuroscience point of view, the results reflect recent findings of computational modelling of scale-free dynamics. Specifically, a large body of literature is showing that computational models of functional connectivity generated from structure show a common pattern across several timescales (Deco et al., 2013, 2009; Ghosh et al., 2008; Hansen et al., 2015; Honey et al., 2007). This speaks to the ICN-FG component and its frequency-independence observed in our study showing equally a common component across all observed timescales.

Conversely, some evidence speaks to additional timescale-specific contributions to functional connectivity organization. Beyond the above-discussed complimentary information that EEG provides when linking structure to function (Wirsich et al., 2017), recent work from Schirner et al. (2018) showed that prediction of empirical fMRI from dMRI can be improved by incorporating EEG data into the modelling. This observation highlights the complimentary information of functional connectivity on several timescales not predicted by the above-described computational models. Indeed, it has been recently demonstrated that tuning the free parameters (such as recurrent connection strength and excitatory subcortical inputs) of a computational model on a local regional level as compared to a global whole brain level helps to better understand the link between the brain structure and brain function (Wang et al., 2019). This local modification of parameters implicates a local signature of dynamics across different timescales on a connection-wise level in line with the second VIS-FS component observed in this study. Indeed, future work could follow this lead and explore the frequency-specific signatures of locally flexible models as proposed by Wang et al. (2019).

### Limitations

Like in any data-driven approach (and in rsfMRI in general) preprocessing steps remain arbitrary. In particular, here the parameters of PCA-based variance reduction and number of final ICs can slightly change the results. We tried to avoid this parameter-dependence by fitting the optimal parameter to the maximum correlation between two independent datasets. Nevertheless, the analysis design still remains flexible and parameter sets most likely also depend on SNR of any given dataset (recording length, field strength of the MR scanner).

### Conclusion

In conclusion, this work sheds new light on the relationship between EEG and fMRI connectivity, suggesting that parts of the joint EEG-fMRI resting-state connectivity is related across timescales and modalities in a spatially independent manner. This data-driven approach based on hybrid extraction of independent connectivity components shows great potential for future research in the field, especially for simultaneous EEG-fMRI in clinical populations.

## Acknoledgements

We thank Katia Lehongre and Benjamin Morillon (main dataset) and Maxime Guye and Jean-Philippe Ranjeva (generalization dataset) for generously sharing their data. JW was supported by a Beckman Institute MoCC seed grant. SS acknowledges financial support from NIH R01MH11622601A1 and R21NS10460302. ALG was supported by ERC 260347 – COMPUSLANG. JG acknowledges financial support from NIH R01EB022574, NIH R01MH108467, Indiana Alcohol Research Center P60AA07611 and the Purdue Discovery Park Data Science Award “Fingerprints of the Human Brain: A Data Science Perspective”.

## Supplementary Information

**Figure SI1:**
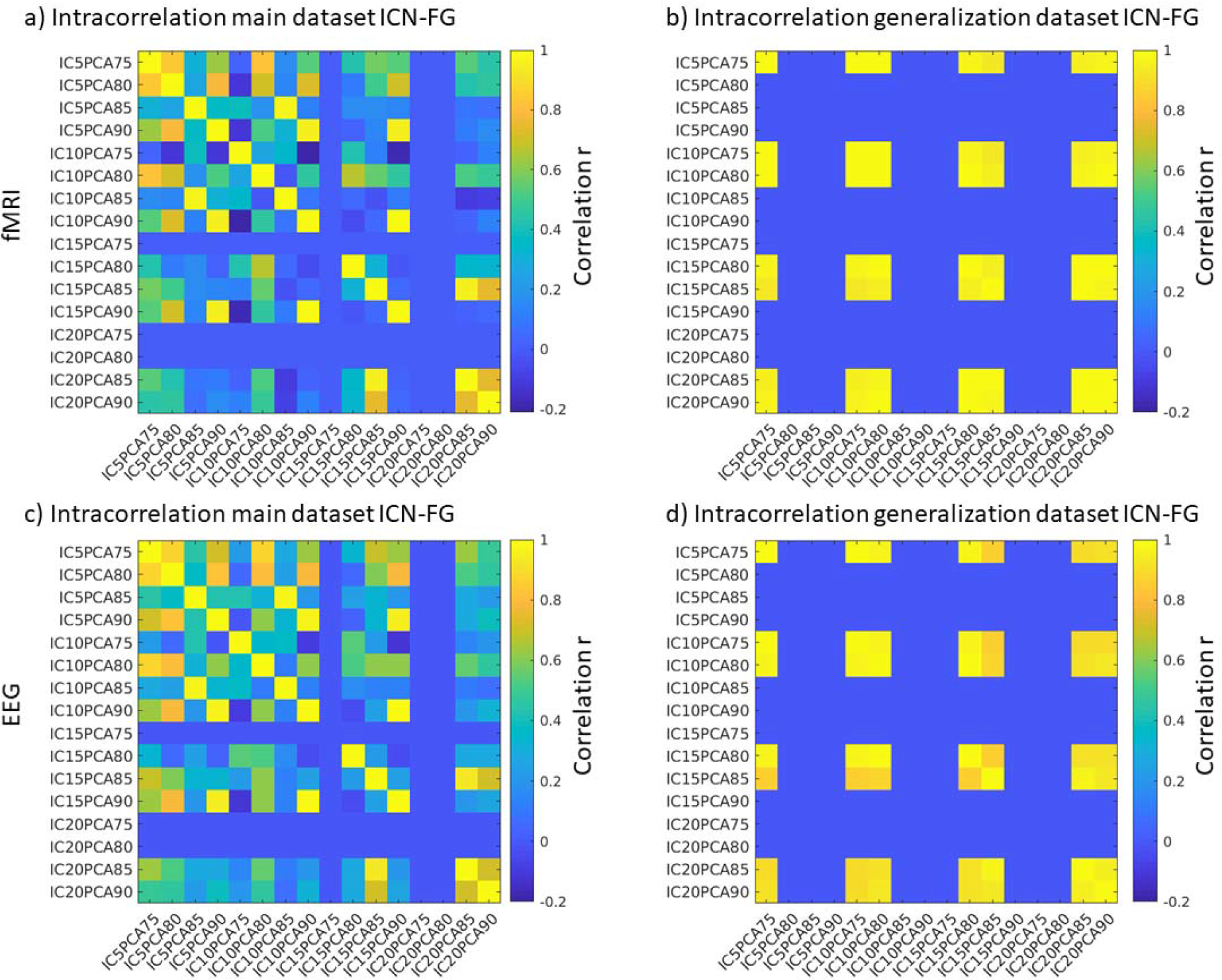
Intracorrelation of the EEG and fMRI part of the IC strengths for ICN-FG component of the main dataset using different parameter sets. We tested all combinations keeping principal components with 75-80% of the variance followed by an independent component analysis keeping 5-20ICs (e.g. IC5PCA75 depicts an analysis using 5ICs and keeping all PCs explaining 75% of the data variance.). Rows with consistent correlation of zero mean that no ICN-FG component was found for this parameter combination.

**Figure SI2:**
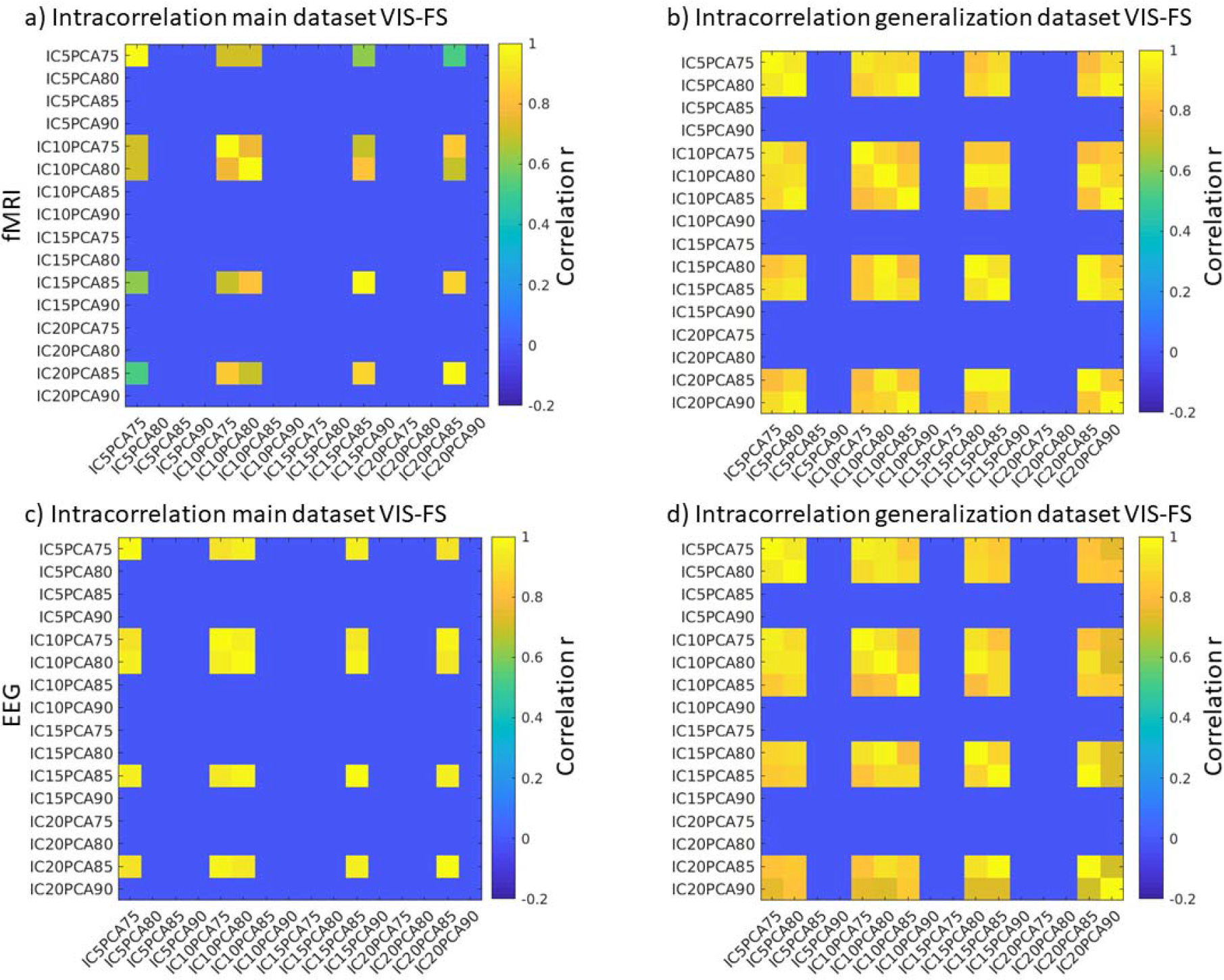
Intracorrelation of the EEG and fMRI part of the IC strengths for VIS-FS component of the main dataset using different parameter sets. We tested all combinations keeping principal components with 75-80% of the variance followed by an independent component analysis keeping 5-20ICs (e.g. IC5PCA75 depicts an analysis using 5ICs and keeping all PCs explaining 75% of the data variance.). Rows with consistent correlation of zero mean that no VIS-FG component was found for this parameter combination.

**Figure SI3:**
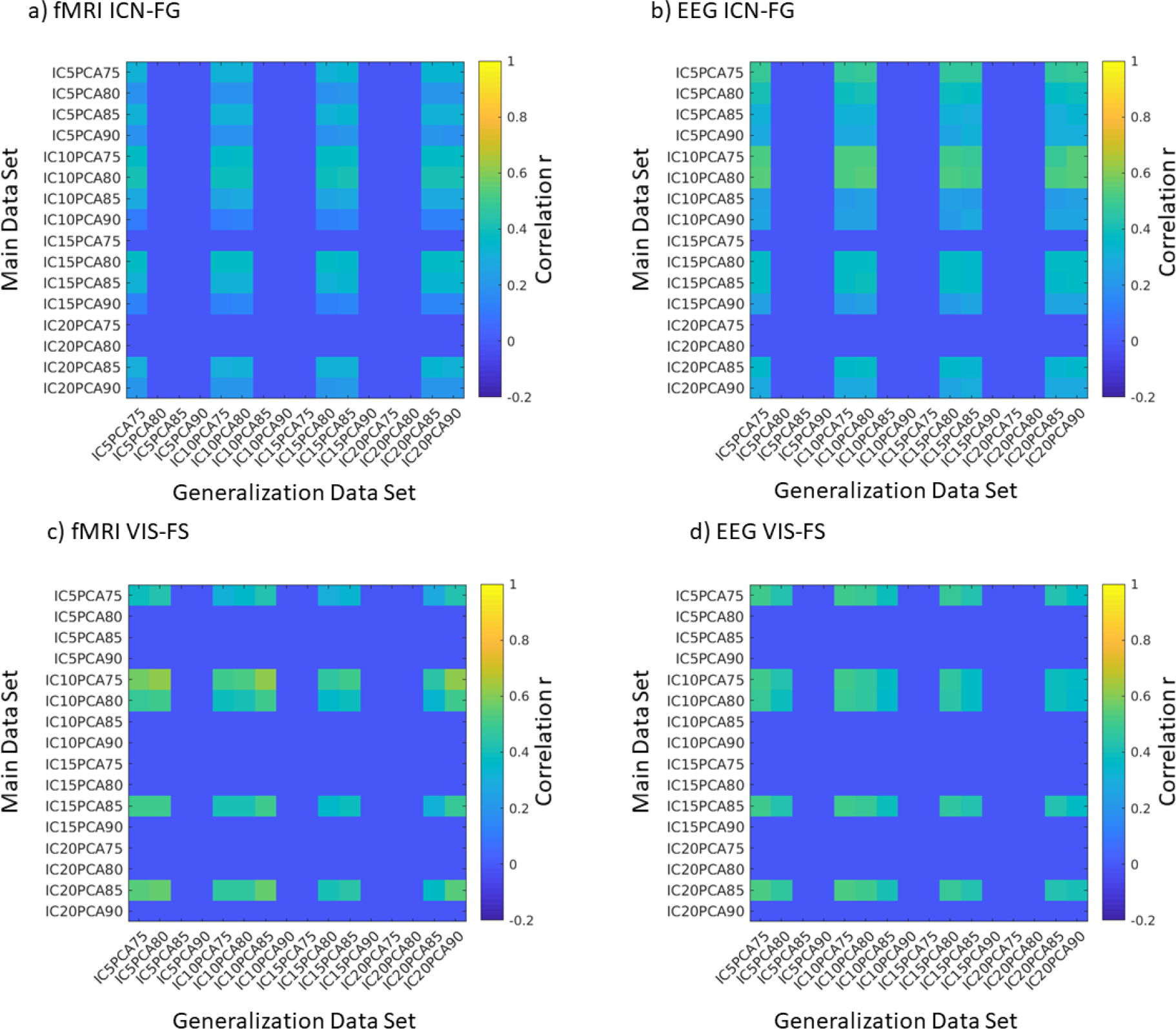
Intercorrelation between both datasets of the EEG and fMRI part of the IC strengths for the ICN-FG and VIS-FS component using different parameter sets. We tested all combinations keeping principal components with 75-80% of the variance followed by an independent component analysis keeping 5-20ICs (e.g. IC5PCA75 depicts an analysis using 5ICs and keeping all PCs explaining 75% of the data variance.). Rows with consistent correlation of zero mean that in one of the datasets no component was found for this parameter combination.

**Figure SI4:**
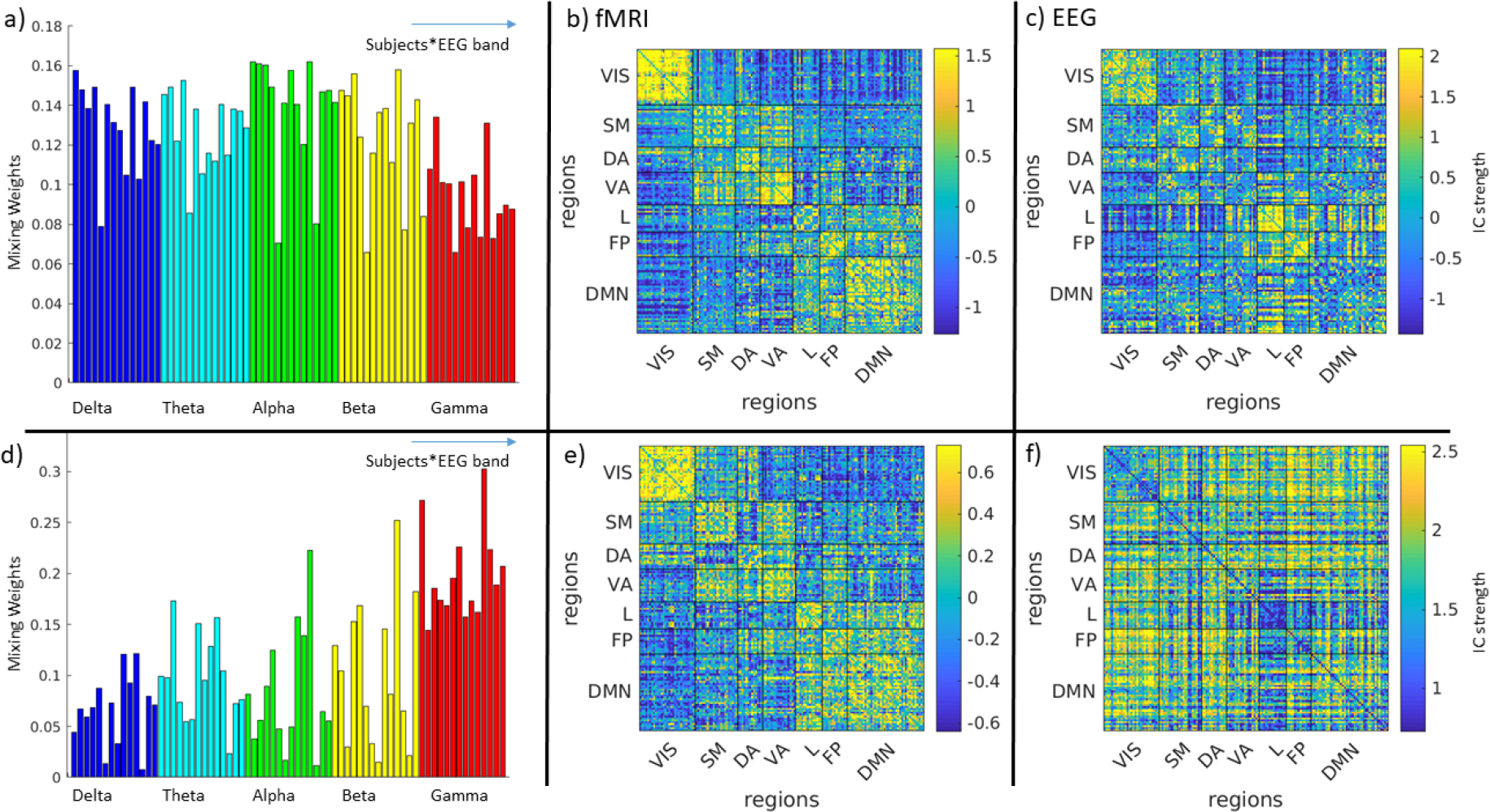
a) subject and band-specific ICA weights of the hybrid EEG-fMRI FG-ICN trait b) fMRI part of the FG-ICN trait c) EG part of the FG-ICN trait d) subject and band-specific IC-weights of the hybrid EEG-fMRI VIS-FS trait e) fMRI part of the VIS-FS trait f) EEG part of the VIS-FS trait. All panels represent the generalization data set (for main data set see Figure 3, colorbars have been saturated at 95^th^ and 5^th^ percentile for better comparison with Figure SI5 and SI6)

**Figure SI5:**
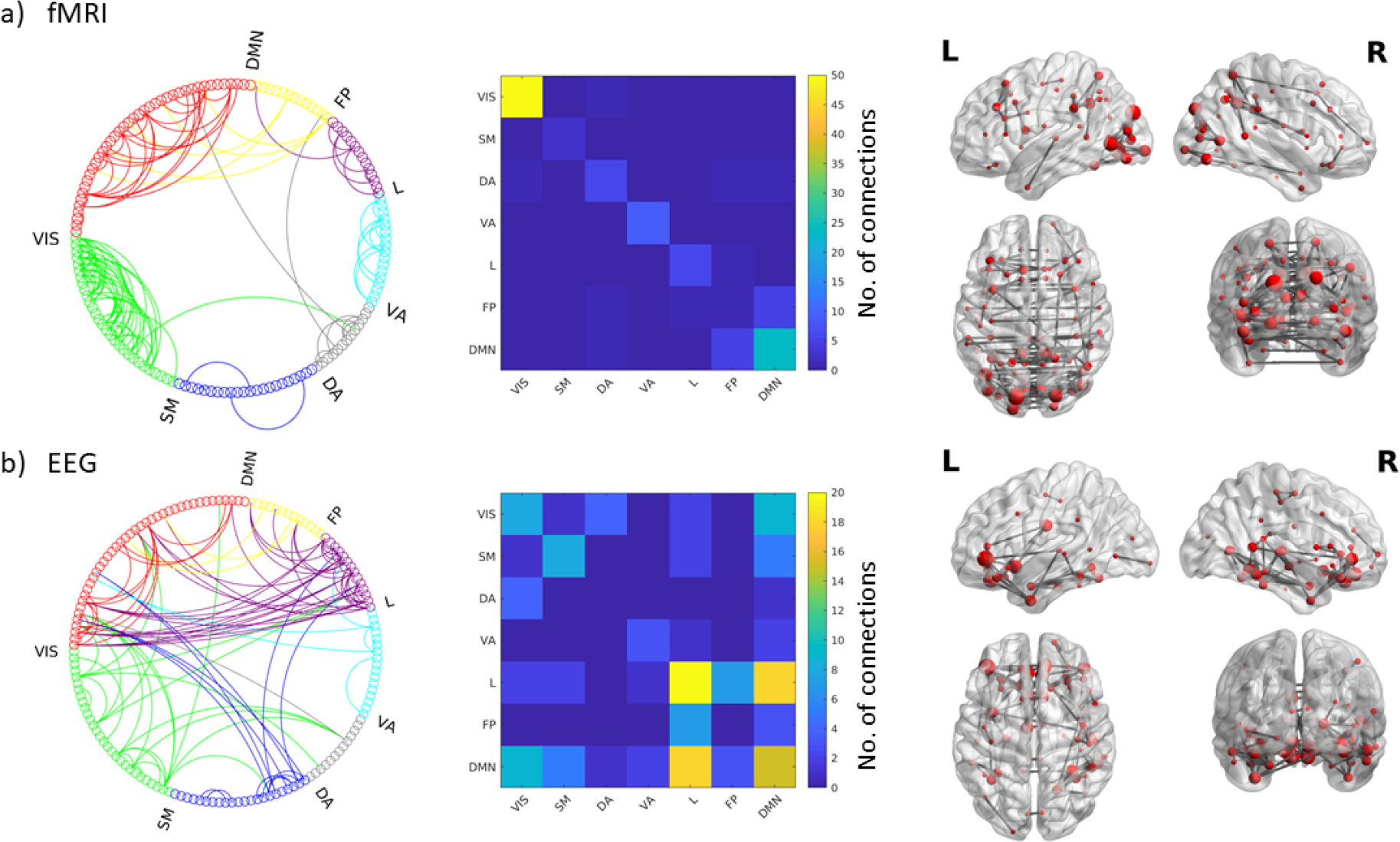
Top nodal strength connections (99^th^ percentile) for the ICN-FG component of the generalization dataset: Circle graphs show all connections between different ICN networks for fMRI (a) and EEG (d); Matrices summarize the number of connections falling into each ICN-ICN pair for fMRI (b) and EEG (e); Brain renderings show strongest connections of the components on a canonical reconstructed cortical surface for the fMRI (c) and EEG (f) part of the hyprid component. Data correspond to the main dataset; for the results if the generalization dataset see Figure SI5. VIS: Visual, SM: Somatomotor, DA: Dorsal Attention, VA: Ventral Attention, L: Limbic, FP: Fronto Parietal, DMN: Default Mode Network

**Figure SI6:**
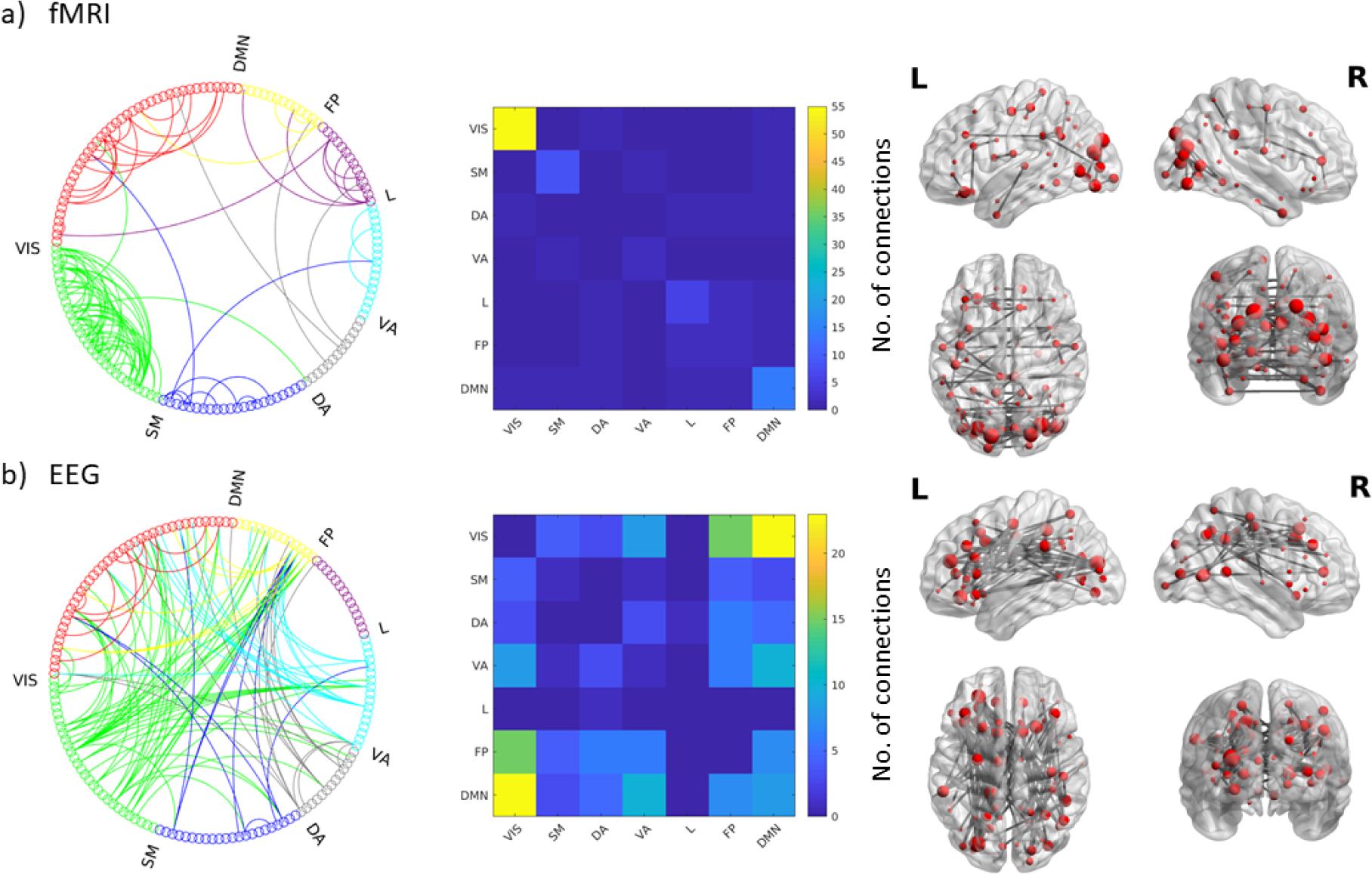
Top nodal strength connections (99^th^ percentile) for the VIS-FS component of the generalization dataset: Circle graphs show all connections between different ICN networks for fMRI (a) and EEG (d); Matrices summarize the number of connections falling into each ICN-ICN pair for fMRI (b) and EEG (e); Brain renderings show strongest connections of the components on a canonical reconstructed cortical surface for the fMRI (c) and EEG (f) part of the hyprid component. Data correspond to the main dataset; for the results if the generalization dataset see Figure SI5. VIS: Visual, SM: Somatomotor, DA: Dorsal Attention, VA: Ventral Attention, L: Limbic, FP: Fronto Parietal, DMN: Default Mode Network

## SI Results

### Supplementary analysis of head motion in the main dataset

As a consequence of the observed correlation between the ICN-FG mixing weights and head motion in the main dataset, we extended the analysis of the main dataset to also look at a previously excluded stable component.

As described in the methods section, the ICN-FG component might split up into two similar components for the main dataset. Indeed, for the chosen parameter set (keeping PCs that explain 75% of the variance and calculating the ICA for 10 ICs), we found a second IC with similar properties to the ICN-FG component namely ICN-organization for EEG and fMRI part of the IC (q_fMRI_ = 0.11, q_EEG_ =0.06), subject specific fingerprint (ICC_subject_ =0.85, p<10^−10^) and no differences of mixing weights as a function of EEG frequency band (ICC_freq_=−0.018, p=0.71). A very small relationship between EEG and fMRI part of the IC weights was observed (r=0.049116, p=2.9722 10^−07^). The IC strengths were associated between the above-described second ICN-FG and the ICN-FG in the generalization dataset (fMRI-fMRI 0.25 p<1.0*10^−300^; EEG-EEG = 0.42, p<1.0*10^−300^). This correlation between main and generalization dataset was lower than the originally found ICN-FG component (both for EEG and fMRI).

Contrasting the observation for the originally found IC, the mixing weights of this IC were not related to movement (FD vs. mixing weights: rho=−0.12, p=0.18; No. of scrubbed volumes vs. mixing weights: rho=−0.02, p=0.78).

